# Screening of the Pathogen box reveals novel anti blood-feeding compounds

**DOI:** 10.1101/2025.08.25.672114

**Authors:** Dovran Ovezgeldiyev, Rory Doolan, Vanessa Trefzer, Carmel Daunt, Melisa Altındağ, Joyce van Bree, Nicola L. Harris, Tiffany Bouchery

**Author notes:** These authors contributed equally to this work and share first authorship (Dovran Ovezgeldiyev, Rory Doolan).

## Abstract

Soil-transmitted helminth (STH) infections such as *Necator americanus* infect millions globally, and are a major cause of anemia and developmental stunting in low and middle income countries. Blood-feeding hookworms in particular rely on the digestion of host erythrocytes for nutrition and therefore detoxify heme as a byproduct of their parasitism. This dependency on blood feeding and subsequent detoxification renders this pathway as a vulnerable target for therapeutic intervention, particularly as it is the cause of morbidity in those infected. Here we described the continued development and application of a high-throughput *in vitro* assay using the so-called rodent hookworm *Nippostrongylus brasiliensis*, a model that shares key traits with *N. americanus* including blood feeding and hemozoin-like pigment formation. We optimized a fluorescence-based screening cascade to utilise GelGreen as a cost-effective viability stain and screened 400 compounds from the MMV Pathogen Box. Multiple compounds displayed enhanced activity in the presence of blood, suggesting interference with blood-feeding or blood-feeding-induced development. Five hits were selected for further validation, and as proof-of-principle of this screening cascade, all five were well tolerated *in vivo* at low doses in a murine model. This study therefore demonstrates this method can be used as a tractable and biologically relevant screening approach to identify compounds active against blood-feeding nematodes. Future work can further develop such compounds into lead drug candidates, and be leveraged for comparative parasitology approaches to identify pan-anthelmintic drugs.

## 1 Introduction

Soil-transmitted helminth (STH) infections represent a major neglected tropical disease, with 1.5 billion people infected around the world. The causative human-infecting agents are *Ascaris lumbricoides*, whipworms (Trichuris spp.), and hookworms (*Ancylostoma duodenale* and *Necator americanus*) which are grouped together as they often respond to the same treatments and water, sanitation and hygiene interventions. Hookworms in particular cause chronic intestinal blood loss and anemia due to the hematophagous nature of the parasite, which burdens adolescent girls and women of reproductive age with morbidity and contributes to the cycle of poverty in low-income countries. As only two treatments, albendazole and mebendazole are readily available for mass drug administration programs, the WHO NTD Roadmap 2030 has called for more effective medicines and drug combinations to be developed against hookworm.

Conventional *in vitro* screening approaches for anthelmintic compounds have largely relied on manual scoring of parasite motility as a readout of viability by light microscopy. While these methods are relatively inexpensive, they are low-throughput, subjective, and require trained personnel to recognise *in vitro* phenotypes and assign categorical scores. Moreover, motility alone may not capture sublethal or subtle developmental effects of drugs, leading to false negatives. Overall, this limits their scalability for large compound library screening. As a result, there is a growing move towards medium-to high-throughput screening methods in the helminth drug discovery field. These range from automated assessments of motility, egg hatching, or ATP-production, and utilise readouts including imaging, electrical impedance, or fluorescence (Dilrukshi Herath et al., 2022).

We have previously described that a strongylid of rodents *Nippostrongylus brasiliensis* shares key features of the blood feeding/heme-detoxification pathway with *N. americanus*, despite being phylogenetically distant to this major human hookworm (Marchand et al., 2022). In particular, we observed a novel pathway of heme detoxification, in addition to glutathione-*S*-transferase, via the production of a hemozoin-like pigment. In line with this, the quinoline compounds quinine, chloroquine, and quinidine were active against *N. brasiliensis in vitro* and *in vivo* and prevented the formation of this hemozoin-like pigment (Bouchery et al., 2018). Here we describe the optimization of a high-throughput drug-screening assay targeting the blood-feeding pathway of *N. brasiliensis*. GelGreen was identified as a cost-effective and robust viability stain for use in fluorescence-based drug assays. We hypothesized that we could use this method with blood-supplemented media to identify novel compounds targeting the blood feeding pathway or parasite development. We therefore screened the MMV Pathogen box, a library of 400 molecules active against neglected diseases. As proof-of-principle we further validated these compounds *in vitro* and *in vivo*.

## 2 Results

### 2.1 GelGreen is a cost-effective and suitable alternative to SYTOX Green for high-throughput screening

Previously, our HTS assay for soil-transmitted helminth hit identification relied on measuring SYTOX Green, a cell impermeant DNA binding dye which stained dead parasite cells. SYTOX Green is costly, and the quality control Z-scores we calculated with this were typically in the “moderate” range (Zhang et al., 1999, Zhang, 2007). We therefore benchmarked SYTOX Green against four other viability dyes: Image-iT DEAD Green, PrestoBlue, GelGreen, and propidium iodide (PI) (Table 1). We incubated healthy, quinidine-treated, or heat-killed larvae with each dye for 3 days to select optimal dye concentrations that could distinguish parasite viability (Supp. Figure 1A). Propidium Iodide, Image-iT DEAD Green, and Gel Green were all able to discriminate between live and dead parasite. In contrary, PrestoBlue could not. Next, we repeated the assay in the presence of blood for 3 days, with or without quinidine, and plate-reader quantification was used to assess staining with the dyes. Image-iT DEAD, PI, and GelGreen all led to a difference in fluorescence of healthy versus quinidine-treated larvae (Fig. 1A). However, Presto Blue failed to distinguish live and dead parasites and was excluded from further testing. Specific staining of parasites was confirmed by fluorescence microscopy (Fig. 1B). While PI showed a strong signal, we detected a high level of background fluorescence (Fig. 1C). We next compared the assay quality with each dye by plating 32 replicates of larvae, and calculating two different quality scores, Z-score and SSMD (Table 1). Altogether, Image-iT DEAD and GelGreen performed well in all aspects we investigated, however considering the price per plate of the assay, GelGreen emerged as a clearly superior choice of dye for our HTS.

**Table 1:** Comparison of viability markers used for drug screening assay.

|  | Details |  |  | Comparison of QND vs DMSO |  |  |  |
| --- | --- | --- | --- | --- | --- | --- | --- |
| Dye concentration | Type | Ex/Em | Top (T) or bottom (B) read | p value | Z' score | SSMD | Price / plate |
| **SYTOX Green 1:100** | DNA | 485/520 | B | <0.001 | 0.47<br>Moderate | 7.675<br>High | 291.0 CHF |
| **Image-iT DEAD Green 1:100** | DNA | 485/520 | B | <0.0001 | 0.30<br>Moderate | 3.269<br>Excellent | 254.2 CHF |
| **PrestoBlue 1:10** | Reduction | 560/590 | T | - | - | - | - |
| **GelGreen 1:500** | DNA | 485/535 | B | <0.0001 | 0.95<br>Excellent | 1.048<br>Moderate | 5.96 CHF |
| **PI 1:10** | DNA | 355/630 | B | <0.001 | 1.71<br>Excellent | 1.075<br>Moderate | 22.0 CHF |

**Figure 1:**
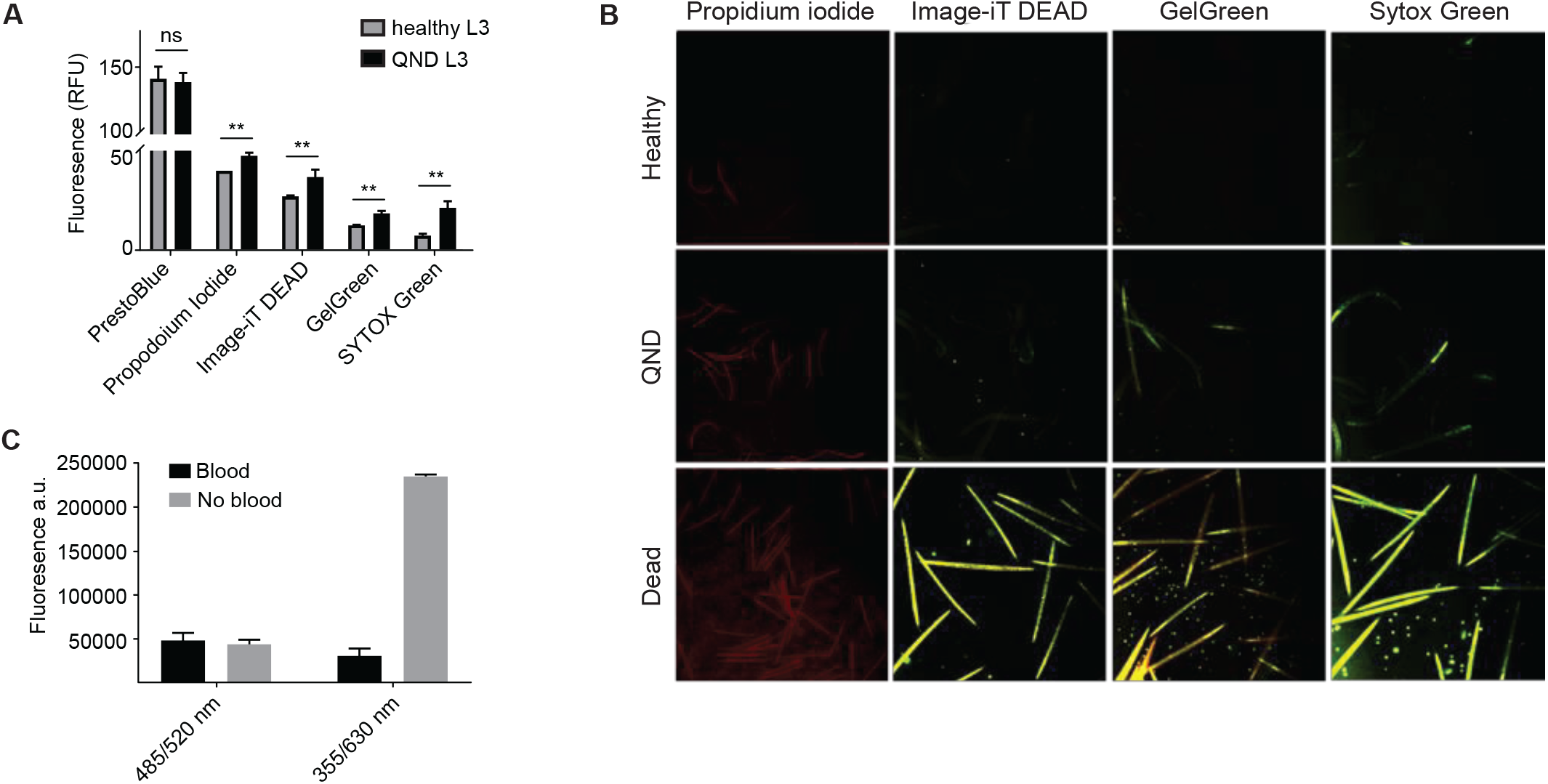
GelGreen is a suitable and cost-effective dye for assessing *N. brasiliensis* L3 viability in blood-supplemented media. Live, dead and quinidine (QND) treated larvae were cultured *in vitro* in the presence of red blood cells. SYTOX Green was added to a final concentration of 1:100, and different concentrations of PrestoBlue, Image-iT DEAD, and GelGreen were added to the larvae after 3 days of incubation. **(A)** Fluorescence was measured at day 4 using an excitation/emission of 485/520 nm and bottom read for SytoxGreen, Image-iT DEAD and GelGreen; 355/630 nm and bottom read for Propidium Iodide; and 535-560/590-615 nm and top read for PrestoBlue. Only the concentrations showing the greatest differences (PrestoBlue 1:10, Propidium Iodide 1:10, Image-iT DEAD 1:100, GelGreen 1:1000) are presented. The data represents 6 replicates of approximately 40 iL3 larvae, Mann-Whitney (nonparametric) test: n.s.: *p*-value > 0.05, ^**^: *p*-value < 0.01. **(B)** Fluorescent microscopy was performed to give a visual representation of the fluorescence of SYTOX Green, Image-iT DEAD, and GelGreen using a blue filter, and Propidium Iodide using a red filter. **(C)** Healthy iL3 larvae were plated with and without blood to check for the effect of hemoglobin on the emission and/or excitation of the different dyes.

### 2.2 High throughput screening of the MMV Pathogen box reveals many blood-feeding hits

Next, we implemented HTS of the Medicines for Malaria Venture (MMV) Pathogen box. Adapting our previous method (Marchand et al., 2022), we cultured L3 with the 400 compounds in the Pathogen box at 100uM for 4 days. HTS was performed using lysed human RBC-supplemented media in order to trigger parasite development and identify blood-dependent larvicidal hits. All compounds were screened twice using two independent life cycles of parasites, and hits were then checked manually by microscopy to assess motility and hemozoin-like pigmentation to rule out false positives from inherently fluorescent compounds. We identified 133 compounds as hits from either screening, 54 of which passed a cutoff (DMSO mean + 2^*^ DMSO SD) in both screenings (Figure 2A & 2B). Microscopic evaluation of larval motility and pigmentation narrowed down the hits to 28 compounds (Table 2). Of these 28, two were reference compounds: the anti-malarial MMV000016 Mefloquine with known pan-anthelmintic properties (Keiser et al., 2009, Lundström-Stadelmann et al., 2020); and MMV000063 sitamaquine a compound under development for visceral leishmaniasis which has shown nephrotoxicity in phase 2 trials (Loiseau et al., 2011, Dietze et al., 2001). Other reference compounds, such as Mebendazole, met the fluorescence threshold but not the phenotypic scoring as they reduced pigment or motility but not both. We further screened all 28 hits in the absence of blood. We found that 7 of these 28 compounds depended on blood-supplemented media for activity, suggesting they target blood-feeding induced development or blood feeding itself (Figure 2C, Table 2). Of these, 4 belonged to the *‘Malaria’* category of the Pathogen Box, and one each to the *‘Tuberculosis’, ‘Cryptosporidiosis’* and the *‘Onchocerciasis’* categories (Table 2).

**Table 2:** List of 28 hits from the MMV Pathogen box screen against *N. brasiliensis* L3. Refined by microscopic scoring from 54 fluorescence-based hits. Compounds marked ^*^ are blood-dependent as per Table 3.

| Compounds | Potency to QND | Microscopic scoring | Category |
| --- | --- | --- | --- |
| MMV688934 | 9.3 | 5 | Kinetoplastids |
| MMV676604 | 5.2 | 5 | Kinetoplastids |
| MMV652003 | 4.9 | 5 | Kinetoplastids |
| MMV021013 | 4.9 | 5 | Tuberculosis |
| MMV675994 | 4.8 | 5 | Cryptosporidiosis |
| MMV688938 | 2.7 | 4 | Tuberculosis |
| MMV062221 | 2.3 | 4 | Malaria |
| MMV637229 | 2 | 5 | Trichuriasis |
| * MMV007920 | 1.8 | 4 | Malaria |
| MMV676401 | 1.6 | 5 | Tuberculosis |
| MMV004168 | 1.6 | 4 | Kinteoplastids |
| MMV000016 | 1.4 | 5 | Reference compounds |
| MMV658993 | 1.4 | 5 | Kinetoplastids |
| MMV000063 | 1.3 | 3 | Reference compounds |
| MMV676512 | 1.3 | 4 | Tuberculosis |
| MMV1029203 | 1.2 | 5 | Malaria |
| * MMV688845 | 1.2 | 4 | Tuberculosis |
| MMV006901 | 1.2 | 4 | Malaria |
| * MMV011511 | 1.2 | 4 | Malaria |
| * MMV001493<br>(Isradipine) | 1.1 | 4 | Onchocerciasis |
| * MMV085071 | 1.1 | 4 | Malaria |
| * MMV026490 | 1 | 4 | Malaria |
| * MMV689255<br>(D-eritadenine) | 1 | 3 | Cryptosporidiosis |
| MMV676602 | 1 | 4 | Kinetoplastids |
| MMV688279 | 1 | 4 | Kinetoplastids |
| MMV020391 | 0.9 | 3 | Malaria |
| MMV090930 | 0.8 | 3 | Tuberculosis |
| MMV688122 | 0.7 | 5 | Tuberculosis |

**Table 3:**
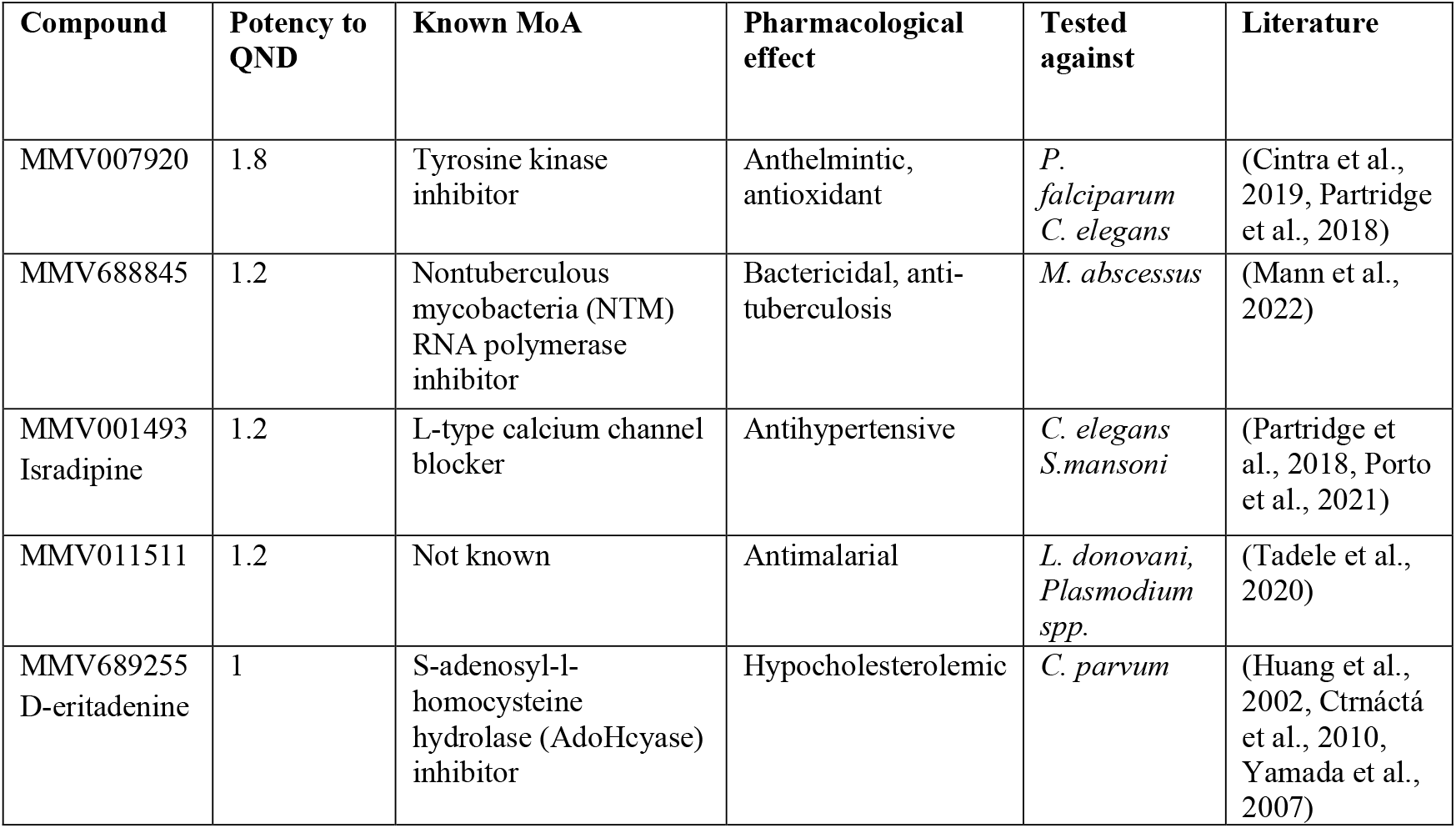
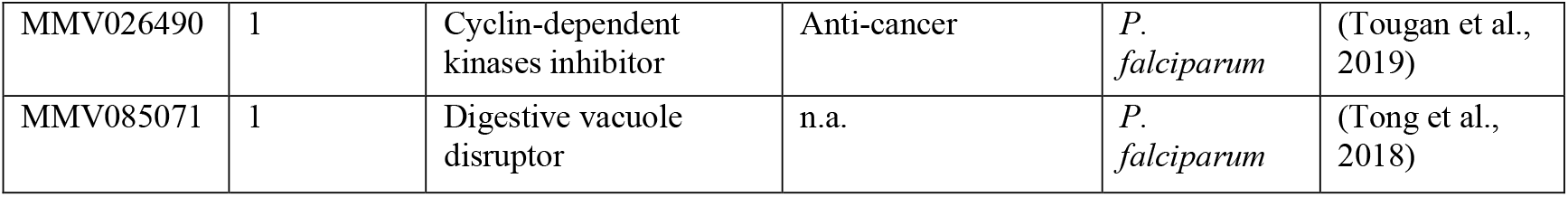
List of blood-dependent MMV Pathogen box compounds with activity against *N. brasiliensis*.

**Figure 2:**
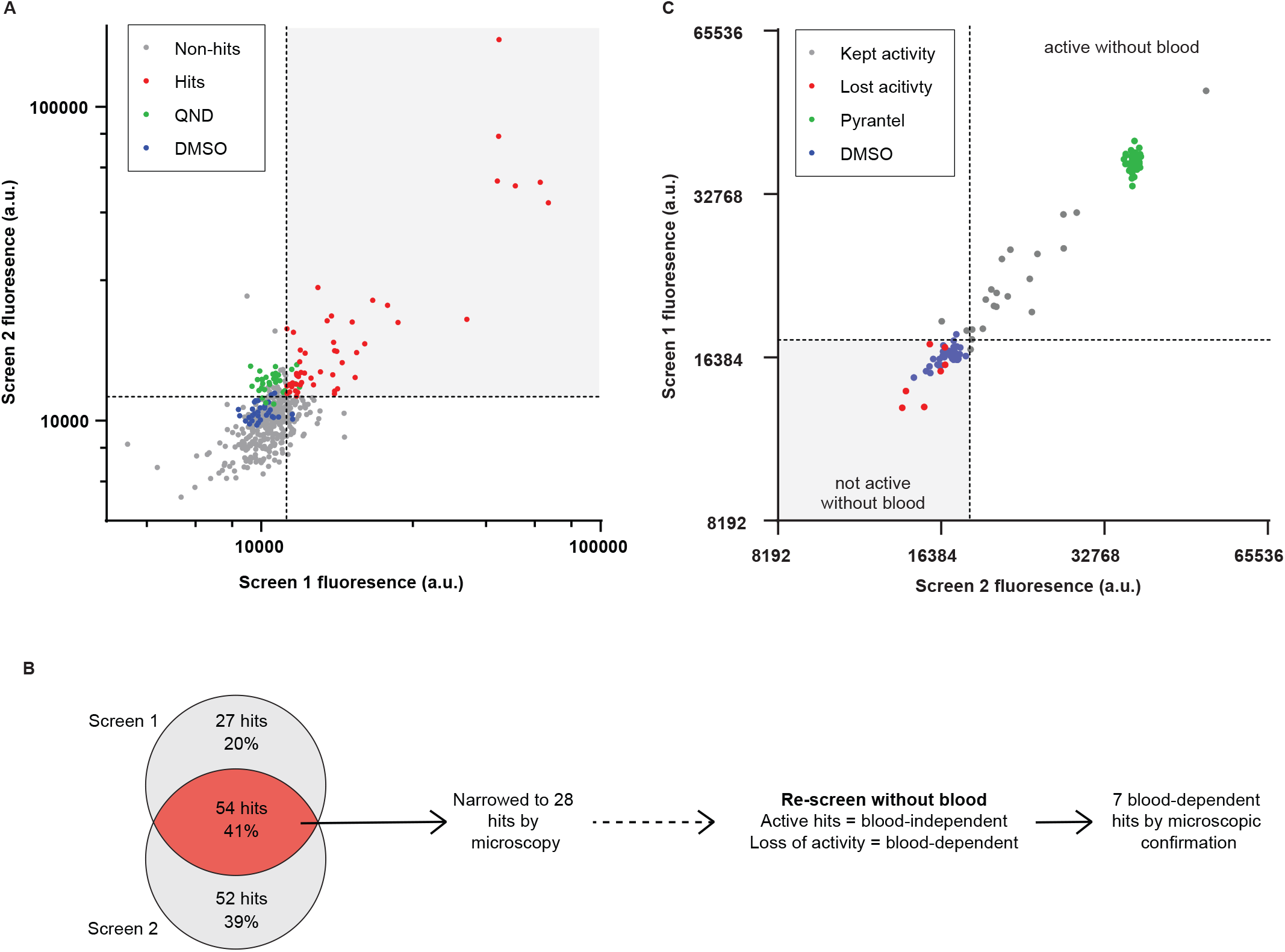
Screening of MMV pathogen box against *N. brasiliensis* L3 in blood-supplemented media revealed 54 hits. Two independent screens of the MMV Pathogen box in the presence of blood were performed. **(A)** L3 were cultured with 100uM of compounds, DMSO, or quinidine (QND) as controls, and the activity threshold was set as 2xSD of DMSO (dotted lines). **(B)** In total 133 compounds were identified as hits across either screen, with 54 double-hits common to both screens. These 54 hits were assessed phenotypically and narrowed down to 28 hits. **(C)** The 28 hits were tested for blood-dependent activity by performing another two independent screens under the same conditions but in the absence of blood. Pyrantel pamoate was used as positive control and the same hit threshold was applied. 7 compounds lost activity in both screens and were assessed phenotypically. Three of these ‘lost activity’ hits were found to be active by phenotypic assessment, and 3 ‘kept activity’ hits were found to not be active by phenotypic assessment, leading to a total of 7 blood-dependent hits being identified from the 28 hits in **(A)**.

Reviewing the literature, 15 of our 28 hits were previously described to be active against other helminth parasite species or *C. elegans*. To leverage our assay for novel drug discovery we next focused on compounds with little to no literature describing anthelmintic activity.

### 2.3 Five anthelmintic compounds of the MMV Pathogen box are safe for further *in vivo* testing

We chose 3 blood-dependent compounds and two potent, blood-independent compounds for further testing. For IC50 determination we used a more traditional manual scoring, which combined pigmentation and motility (Figure 3A). An IC50 could not be calculated for MMV011511 as it is likely above the 100uM range. Absolute IC50s were calculated for the other four compounds: MMV085071, MMV688845, MMV675994, and MMV021013 (Figure 3A). MMV675994 had the lowest IC50 of 1.13uM against *N. brasiliensis* L3. After reviewing MMV cytotoxicity literature, we concluded that all five compounds were suitable to test *in vivo*. To ensure that drug-parasite interactions did not exacerbate adverse events, we tested all five compounds in infected mice and used an oral route of administration as this is an essential characteristic of new drugs for STH (Keiser, 2023). A low dose (5mg/kg) was administered orally for two days on the third and fourth day of infection, during the hemorrhaging phase of lung-to-gut migration. Twice per day we weighed mice and monitored for adverse drug or drug-parasite reactions. No adverse reactions to the compounds were observed, and the body weight change of each drug treated group was not significantly different than the vehicle control group (Fig 3B). Intestinal worm burden was measured 5 days post-infection, and at this low dose of 5mg/kg no drug activity was observed for any of the compounds. Due to limited supply of compounds, it was not possible to test a higher dose *in vivo*.

**Figure 3:**
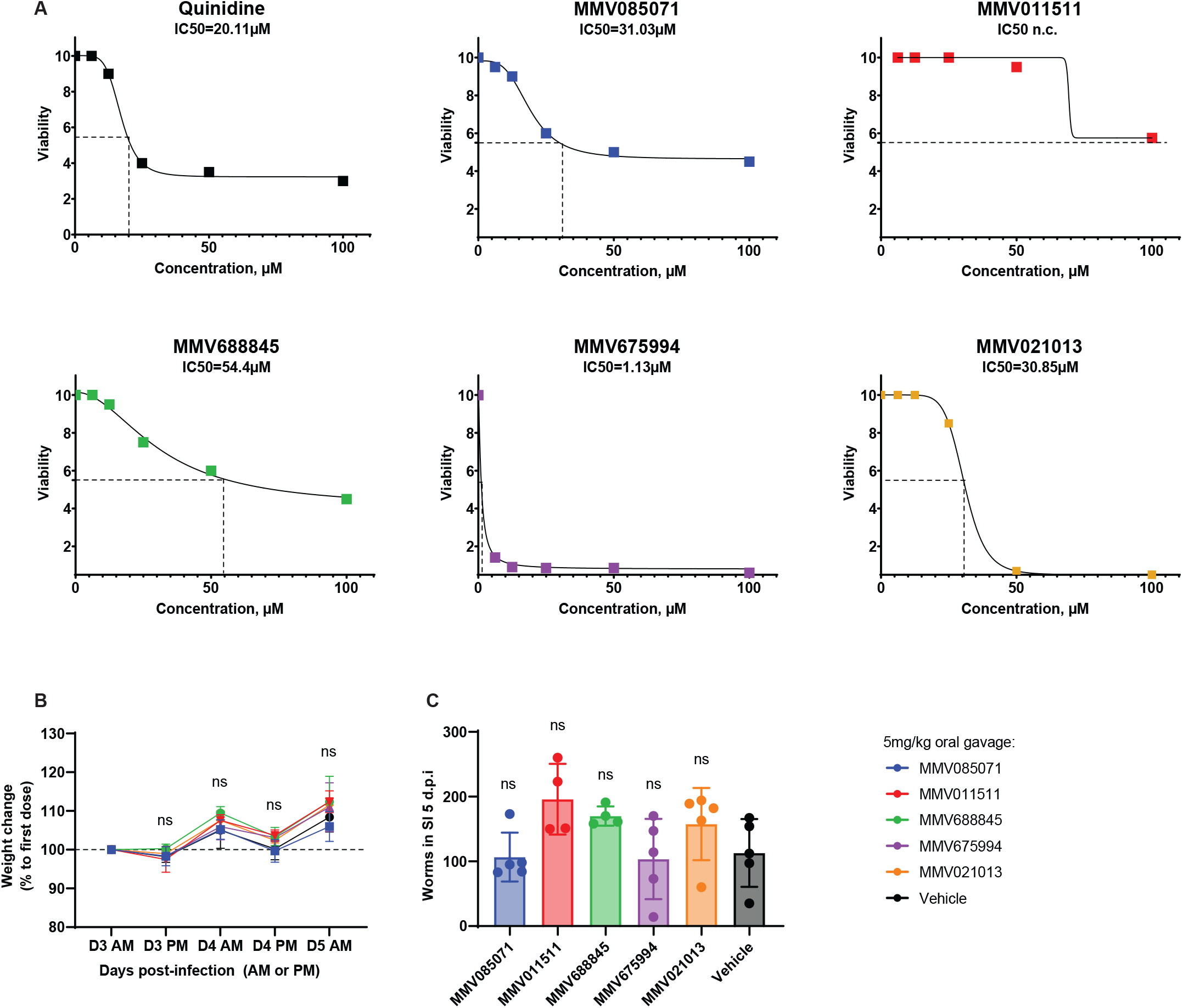
Activity and safety profiles of 5 compounds from the MMV pathogen box. Further testing was performed on 5 hits from HTS screening against *N. brasiliensis*. **(A)** Phenotypic scoring was used for absolute IC50 determination: the frequency of larvae with normal pigmentation and motility (compared to DMSO controls) was scored 1-10, then an average score was calculated to incorporate both phenotypes, with 5.5 corresponding to a combined inhibition of 50%. **(B&C)** Mice were infected with Nb by subcutaneous infection on D0. 3 and 4 days later the compounds were administered orally at 5mg/kg. **(B)** Mouse weight was monitored twice daily (AM or PM) between 3 and 5 days post-infection **(C)** Worm burden was evaluated in the small intestine (SI) by baerman 5 days post-infection (d.p.i). Ordinary one-way ANOVA with Dunnett’s multiple comparisons testing was performed for worm burden data. A mixed effects model was used to assess the % weight change of mice, with Dunett’s multiple comparisons used to compare experimental groups on each day. ns: *p*-value >0.05.

## 3 Discussion

This study provides a proof-of-concept that *N. brasiliensis* L3 larvae cultured with blood can serve as a starting point for a screening cascade to discover anthelmintics which could target blood-feeding or parasite development. In this way, our assay moves closer to a physiologically relevant system by using media supplemented with host blood. GelGreen emerged as a robust and cost effective alternative to SYTOX Green, enabling scalable fluorescence-based HTS. Furthermore, the use of human blood in our follow up assays further refined our method in line with the 3Rs principles.

By incorporating blood into our assay, we found a large number of compounds, 54 hits that met the initial criteria for activity. This number is larger than comparable screening cascades in the literature, for example, when the same library was screened against the sheep parasite *Hemonchus contortus* in blood (and serum)-free conditions, only one hit, Tolfenpyrad, could be identified from the same box (Preston et al., 2016); we also identified this as the most potent of our 28 hits, MMV688934. Improving the physiological relevance of our assay therefore led to many hits being detected. Recently, Thilakarathne et al. (2025) screened the MMV Global Priority Box against *H. contortus* with and without the addition of sheep serum. Using a conventional threshold of motility reduction, they identified 6 compounds active in the absence of serum, but surprisingly only 1 compound active with serum. The authors suggest that this reduction in the number of hits with serum is caused by the development of parasites that begins *in vitro* with serum supplementation, together with a decrease in compound availability due to protein-binding. It is important to note that both our blood-supplemented and our base media contain 10% FCS. Adjusting their threshold, Thilakarathne and colleagues identified 2 (out of 240) compounds that were only active in the presence of serum; both were known anthelmintics. This finding highlights that unique compounds can be discovered using physiologically relevant systems, but importantly that as parasite biology changes with host-adaptation, susceptibility to certain compounds also changes.

Interestingly, when we searched the literature for known modes of action of the 7 blood-dependent hit compounds, we concluded that most of the compounds were probably inhibitors of pathways or enzymes downstream of developmental pathways, rather than inhibitors of heme detoxification specifically. While hemoglobinolysis in hookworms (both *A. caninum* and *N. americanus*) is known to rely on a sequential enzymatic pathway, the subsequent detoxification is less well characterized (Ranjit et al., 2009, Williamson et al., 2004). Glutathione-*S*-transferase, a vaccine candidate, is thought to neutralise toxic byproducts of heme, and hemozoin-like pigment formation is inhibited by quinolines, such as the quinidine used as a control in our screen. Given the limited molecular characterisation of pigment formation, it could be that redundant pathways of heme detoxification exist, therefore limiting our ability to detect compounds that only disrupt one of these hypothetical pathways.

By comparing blood-fed and blood-free drug conditions, we were able to identify 7 compounds that displayed anthelmintic activity in the presence of blood only. In addition to the developmental pathways described above, it is also possible that the addition of lysed red blood cells introduces drug-metabolising enzymes which could generate active metabolites. While phase I metabolism is more commonly associated with inactivation of compounds, RBCs are known to possess a high quantity of esterases, methyl transferases, and deaminases (Dash et al., 2021). Artesunate, for example, is metabolised from its pro-drug form by red blood cell esterases to its active metabolite (Morris et al., 2011). In this way, our blood-dependent hits may be pro-drugs activated by host-derived enzymes. In general, however, the lack of mitochondria, nuclei, and cytochrome enzymes in red blood cells means they have limited drug-metabolising activity. To interrogate the blood-dependency of these drugs, future investigations into these 7 compounds could assess the effect of supplementing the media with blood on different days of the assay, or by performing IC50s with a fixed concentration of each compound but a dilution series of lysed blood concentrations to detect for whether a dose-response relationship exists. However, such a method would require a non-pigmentation-based scoring system.

The three most potent hits in our screen were related to kinetoplastids, however these were not blood-dependent. Here we define blood-dependent as hits that lost activity when re-screened in the absence of blood (Figure 2C). Blood-dependent compounds might target biological pathways related to development, whereas ‘blood-specific’ compounds would disrupt blood feeding leading to the accumulation of toxic heme in the parasite, or deprivation of nutrients normally derived from blood. We have not confirmed this specificity in our assay. The toxicity of such compounds, and likely the intensity of fluorescence signal, is limited by the availability of heme in our assay. Additionally, we confirmed the hits from the plate-reader assay with fluorescence microscopy, and observed that some compounds only induced staining (i.e. cell death/permeability) in discrete regions of the parasite such as the intestinal lining or pharynx region of the head (data not shown). This non-homogenous staining also means that toxicity is not directly related to fluorescence intensity for all compounds, and therefore potency/fluorescence signal should be interpreted with caution. We overcame this limitation by implementing a more traditional motility and pigmentation-based manual scoring system later in the screening cascade.

We detected several reference compounds in our screens, and two made it through our filtering: mefloquine and sitamaquine. Mefloquine, a synthetic analogue of quinine has been shown to have activity against schistosomes in addition to its well-known antimalarial activity (Holtfreter et al., 2011). However, an exploratory clinical trial did not observe any efficacy of mefloquine against STH in school-aged children in Côte d’Ivoire (Keiser et al., 2010). Interestingly, sitamaquine has not to our knowledge been reported to have anthelmintic activity, but nephrotoxicity has already been observed in a clinical trial. Other side effects observed were consistent with those caused by 8-aminoquinoline compounds in patients with G6PD-deficiency, so this compound would likely be incompatible with preventive chemotherapy (Loiseau et al., 2011).

To follow up on the hits identified in the high-throughput screen, we performed IC50 determination using human blood. This switch, necessitated by ethical limitations, may have altered the assay conditions. To address this, we halved the concentration of cells when using human RBCs. Not only does the hemoglobin content of human and mouse RBCs differ, the biochemical and protein composition is also different. Nevertheless, species-specific RBC properties should be considered when using non-host-derived blood. We tested 5 compounds from the MMV Pathogen box *in vivo* in an infection model after reviewing known cytotoxicity information from MMV. ADME data was also available; however this was only measured in plasma which may not be relevant for lumen-dwelling parasites. No adverse reactions were observed at a low dose. Further work should investigate higher dose of those compounds in *in vivo* treatment as well as *in vitro* studies using adult worms.

Altogether, the GelGreen method used here considerably lowered the cost of screening compared to our previously published assay; the use of human RBCs improved the ethical outlook of this screening; and in a proof-of-principle we could implement a screening cascade from library to *in vivo* safety testing.

## 4 Materials and Methods

### 4.1 Animals and ethics

Female Lewis rats were obtained from ARC (Australia) or Charles River (Basel) and maintained at Monash Animal Research Platform (Alfred Alliance campus, Australia) or the Swiss TPH (Allschwil, Switzerland) and used between 6-10 weeks of age. C57BL6/J female mice (Charles River) were used between 8-12 weeks of age. All experiments were approved by and performed in accordance with ‘*The Alfred Medical Research and Education Precinct Animal Ethics Committee*’ under reference E/1893/2019/M, or by Kanton Basel-Landschaft under reference BL/543/2021 and BL/560/2025.

### 4.2 Preparation and isolation of *N. brasiliensis*

The life cycle of *N. brasiliensis* (the Prof. Lindsay Dent strain kindly provided by Prof. Graham LeGros, MIMR, New Zealand) was maintained in rats as described previously (Camberis et al., 2003). In brief, rats infected with 3000-5000 infective third-stage larvae (iL3s) were placed on grids overnight during patency and the feces collected and processed as previously described. iL3s were collected from the edges of coproculture plates for use *in vitro* and *in vivo* within 3 weeks after emergence.

### 4.3 Preparation of blood

Mouse red blood cells (RBC) were collected by cardiac puncture or cheek vein bleeding at euthanasia: for dye optimisation, EDTA tubes (Microvette®200 K3E) were used; for screening assays blood was collected directly into Alsevers solution. RBCs were washed three times in RBC-wash buffer (containing 21.0 mM tris(hydroxymethyl)aminomethane, 4.7 mM KCl, 2.0 mM CaCl_2_, 140.5 mM NaCl, 1.2 mM MgSO_4_, 5.5 mM glucose, and 0.5% bovine albumin fraction V; pH 7.4), resulting in 98% purity (as identified by Ter119-staining by flow cytometry).

Blood-supplemented medium was prepared by lysing RBCs in distilled water and reconstituting in 10X RPMI supplemented with 10% FBS (Gibco), 2 mM L-glutamine (Gibco), 100 U/mL Penicillin/Streptomycin, 0.1 mg/mL Gentamicin). For IC50 experiments, human blood was used after a side-by-side comparison to mouse blood. Blood was provided by the Blutspende SRK Beider Basel blood bank in EDTA K2 tubes, and processed as above. In blood media, mouse or human RBCs were used at a final concentration of 0.5×10^8 RBC/well and filtered using a PES 0.2um filter to remove precipitates.

### 4.4 Preparation of larvae

iL3s were washed three times by gravity sedimentation in phosphate-buffered saline (PBS) and activated for 30 min to 1 h at 37°C in an antibiotic solution (Penicillin/Streptomycin 1000 U/mL (Gibco), Gentamicin 300 U/mL (Sigma) in PBS). For all subsequent steps larvae were kept at 37°C in complete RPMI medium (Gibco) supplemented with 10% FBS (Gibco), 2 mM L-glutamine (Gibco), 100 U/mL Penicillin/Streptomycin, 0.1 mg/mL Gentamicin).

### 4.5 Handling of drug compounds

MMV Pathogen box was stored at –80 °C in 96-well plates. Intermediate stocks (1 mM in RPMI) were made and stored at –80 °C. High-throughput screening was performed at a final concentration of 100uM, with 100uM of Quinidine (Sigma-Aldrich, Merck), 100uM of pyrantel pamoate (Sigma-Aldrich) or 1% (v/v) DMSO (PanReacAppliChem, ITW reagents) added to control wells.

### 4.6 *In vitro* culture and screening assay

Larvae were co-cultured in 30 µL of medium (20uL blood medium + 10uL or larvae medium) in 384 well plates (Greiner bio-one, Fischer Scientific), with 40 iL3 per well, and compounds at 100 µM for 4 days at 37 °C, 5% CO_2_. On the third day, either SYTOX Green™ (Invitrogen™), Propidium Iodide (stock: 2 mg/mL; AnaSpec), Image-iT DEAD Green™ (ThermoFisher), PrestoBlue™ Cell Viability Reagent (ThermoFisher), and GelGreen® Nucleic Acid Gel Stain (Biotium) were added at the concentration mentioned in the figure. For HTS, GelGreen was used at a dilution of 1:500 (from the 10000X commercial stock, stock in DMSO). Spectrophotometric measurements were taken 24 h later using an automated microplate fluorometer (FLUOstar Omega, BMG Labtech, or Spark, TECAN) using appropriate excitation/emission measurements (Table 1).

### 4.7 Microscopic scoring of larvae

Phenotypic screening of motility was performed on hits from the HTS assay to confirm localization of the fluorescence signal to larvae (*i*.*e*. exclusion of false positives due to drug autofluoresence or contaminants). For IC50 determination, larvae were scored microscopically by assessing the % of larvae per well with pigmentation, and the % of motile larvae. An average was taken and scaled from 1-10 (i.e. ‘1’=0-10%… to ‘10’=91-100%); therefore 1 = unhealthy larvae, 10 = healthy larvae, and 5.5 corresponds to an average of 50% inhibition for absolute IC50 determination.

### 4.8 Drug tests *in vivo*

C57BL6/J female mice (8 weeks of age) were ordered and allowed to acclimatize for two weeks after arrival (temperature approx. 25 °C; humidity approx. 70%; 12-hour light and 12-hour dark cycle) with access to water and rodent chow *ad libitum*. Mice were housed in individually ventilated cages (5 per cage) in a conventional facility and infected with 500 *N. brasiliensis* L3 in 100uL phosphate-buffered saline (PBS) by subcutaneous injection in the neck using a 25G needle (D0). In the morning of day 3 (D3) of infection, a single dose of 5mg/kg of the specified MMV compound (approx. 50uL) diluted in DMSO was administered by oral gavage using a round-tipped gavage needle. A second dose was repeated on D4. Detailed welfare monitoring was performed twice daily, and mice were weighed in the morning and afternoon. On D5 of infection, mice were euthanized by CO2 inhalation and the lungs, stomach, and intestines were dissected. Minced tissues were suspended in cheesecloth in a 50mL Falcon tube Baermann apparatus with room temperature PBS, and placed in a water bath which was then set to 37°C. At least two hours later, sedimented worms were collected and counted under a dissecting microscope.

### 4.9 Statistical methods

Z’ score and SSMD were calculated using the formulas:

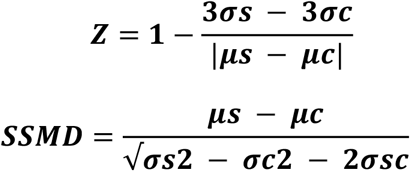

The hit selection cut-off was calculated using:

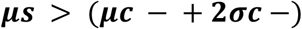

Worm burdens of drug-treated groups were compared using one-way ANOVA with Dunnett’s multiple comparisons.

## 6 Figure Legends

**Supplemental Figure 1:**
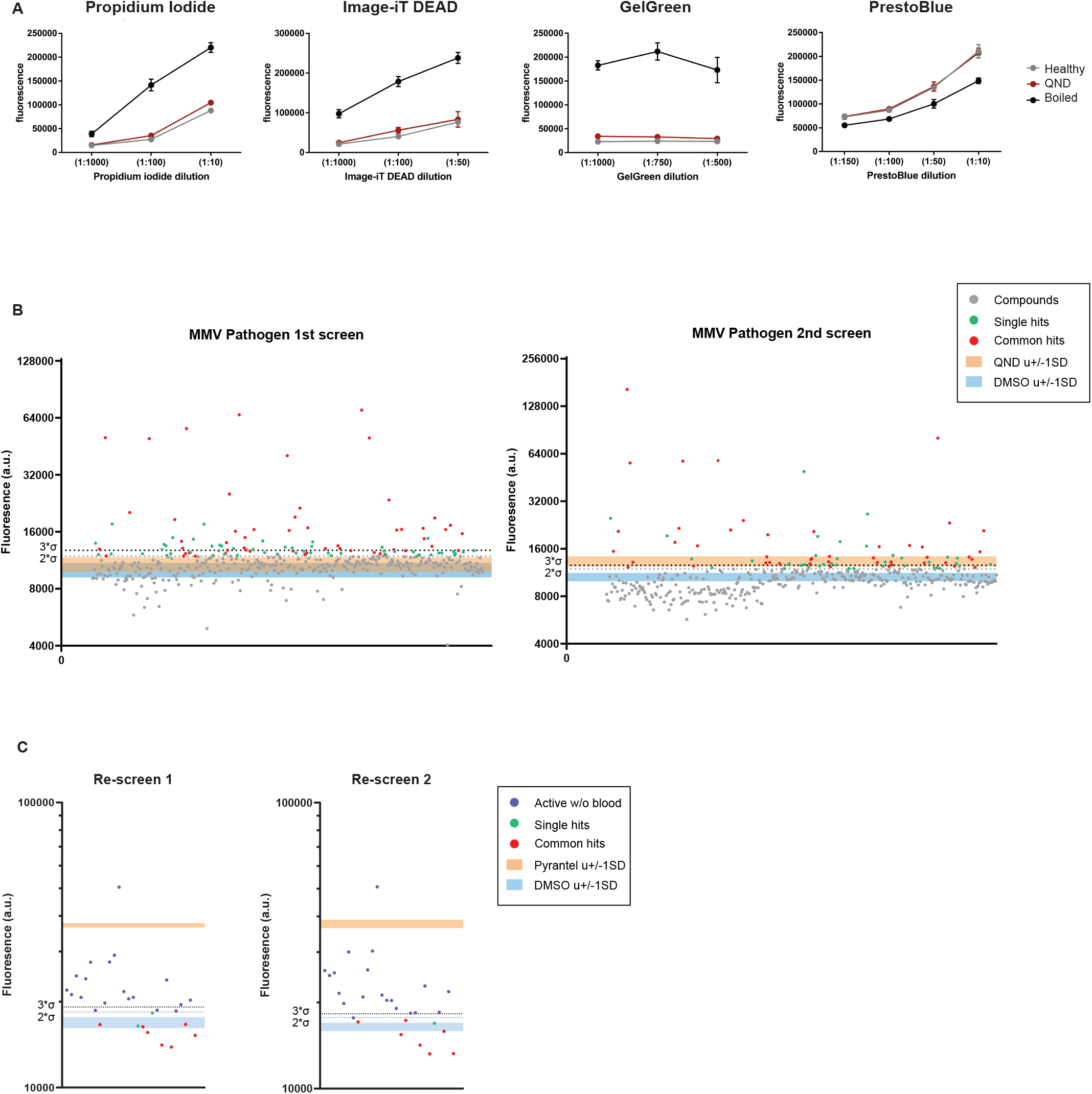
Screening of MMV pathogen box against *N. brasiliensis* L3 in blood-supplemented media. Drug screening results as in Figure 2 as independent screens. Orange and Blue bands represent the mean +/-SD of QND or DMSO respectively, or Pyrantel pamoate or DMSO respectively. Dotted lines represent DMSO mean +2xSD or +3xSD; only the ‘2xSD’ was used for hit identification.

## 7 Conflict of Interest

*The authors declare that the research was conducted in the absence of any commercial or financial relationships that could be construed as a potential conflict of interest*.

## 8 Author Contributions

DO, RD, designed and performed the experiments. JVB performed experiments related with the dye optimisation. VT and CD maintained the parasite life cycle and supported with experiments. MA performed *in vivo* experiments. TB and NH conceived the project and obtained funding. All authors participated in the writing of the paper.

## 9 Funding

This research was funded by Swiss National Science Foundation (grant number PR00P3_193084), European Society of Clinical Microbiology and Infectious Diseases (grant number Research Grant 2017), National Health and Medical Research Council (grant number NHMRC SRF-B), Australian Research Council (DP210101500).

## 10 Acknowledgments

The authors thank Dr. Sarah Preston for sharing her valuable experience and feedback with us. We are grateful to MMV for their support and for designing and supplying us with the MMV Pathogen Box and additional compounds for further testing.

## 11 Data Availability Statement

The datasets generated for this study can be found in the supplementary datasheets.

## Notes

### Competing Interest Statement

The authors have declared no competing interest.

